# Praja1 protects cells from DNA damage through direct DNA binding

**DOI:** 10.64898/2025.12.04.691747

**Authors:** Kotaro Kawasaki, Toru Asahi, Wataru Onodera

## Abstract

Praja1 is known as an E3 ubiquitin ligase that regulates multiple functions through protein degradation. It acquired nuclear localization signal after gene duplication and although studies have shown some significant roles of nuclear Praja1, comprehensive analysis still lacks. In this study, we performed comparative proteomics and biochemical analyses to elucidate the functions of Praja1 in the nucleus. First, proteomics analysis applied to nuclear localization deficient Praja1 exhibited signs of DNA damage response fluctuation. Subsequent comet assay revealed Praja1 protecting cells from various DNA damage sources. Similarly, cells lacking Praja1 became more sensitive to DNA damage-induced cell death, while *E. coli* expressing Praja1 exhibited resistance to DNA damage. To further elucidate the molecular basis of DNA protection, gel shift assay showed direct binding of Praja1 to DNA through electrostatic interactions within its intrinsically disordered region. Further in vitro damaging assay suggested that Praja1 may induce structural changes that enhance DNA repair efficiency upon binding to DNA. Together, these results provide insights into the evolutionarily novel role of nuclear Praja1 in protecting against DNA damage.

## Introduction

Praja1 is an E3 ubiquitin ligase that promotes proteasome-dependent degradation of specific substrates through their recognition and ubiquitination. Depending on its target substrates, Praja1 regulates diverse cellular processes including cell differentiation, cell cycle progression, and apoptosis [1-3]. In the nervous system, accumulating evidence suggests that Praja1 is involved in both neurodegenerative disease pathogenesis and normal neural development [4-6]. For instance, reduced Praja1 mRNA levels have been observed in brain tissues from Alzheimer’s disease patients, while decreased protein levels are found in the 5xFAD mouse model [7,8]. Furthermore, the microtubule-associated protein tau, a major component of neurofibrillary tangles in AD, and spinophilin, which is closely associated with disease progression, have been identified as Praja1 substrates [9]. In addition, a loss-of-function Praja1 mutation (R376C) has been reported in Japanese patients with neurodevelopmental disorders, indicating that Praja1 plays a crucial role in normal nervous system development [10-11].

Our molecular evolutionary analysis revealed that Praja1 arose through gene duplication in the common ancestor of placental mammals [12]. Importantly, Praja1 acquired a nuclear localization signal (NLS) after this duplication event, enabling its nuclear localization. This functional innovation may have allowed Praja1 to recognize and degrade a novel repertoire of nuclear substrates. For example, MSX2, a transcription factor, is a nuclear substrate of Praja1, and MSX2 mutants found in Boston-type craniosynostosis patients undergo excessive ubiquitination and degradation by Praja1 [13]. In satellite cells and C2C12 cells, nuclear Praja1 promotes normal myogenic differentiation by degrading the transcription factor EZH2 [14]. While these nuclear substrates demonstrate that Praja1 functions within the nucleus, the full spectrum of nuclear Praja1 functions remains incompletely characterized.

In this study, we discovered that nuclear Praja1 exerts a DNA-protective function through direct interaction with DNA. Initially, comparative proteomic analysis using NLS-deleted Praja1 revealed that nuclear Praja1 associates with molecules involved in the DNA damage response pathway. Indeed, Praja1 expression reduced DNA damage upon its induction, whereas Praja1 knockdown increased both DNA damage and cell death, suggesting a DNA-protective role for Praja1. Remarkably, Praja1 also suppressed DNA damage-induced cell death in *E. coli*, which lack endogenous Praja1-interacting partners, indicating that Praja1 can interact with DNA independently of other molecular partners. To elucidate the molecular basis of this DNA-protective activity, we examined the Praja1-DNA interaction and found through gel shift assays that the extensive intrinsically disordered region (IDR) of Praja1 interacts with DNA via electrostatic interactions. Further in vitro study suggested that Praja1 may directly induce structural changes that facilitate efficient genomic DNA repair.

## Materials and methods

### Protein purification

Praja1 protein purification was performed essentially as previously described [9]. Briefly, pET-HisTEV Praja1 vector or ΔRING Praja1 vector lacking the RING domain was transformed into Rosetta2 (DE3) cells (#71397, Merck Millipore). This mutant was produced by using forward primer: 5’-TAAGAATCTATTGATGCCCTGCCGG-3’ and reverse primer: 5’-AGATGCCGGAGGATTGGCAACCTC-3’ utilizing KOD -Plus-Mutagenesis Kit (TOYOBO, #SMK-101). After adding 5 mL of medium to the plate and suspending all colonies, cells were collected in 250 mL of 2×YT medium and cultured at 37°C with shaking at 170 rpm until OD_600_ reached 0.5–0.7. For induction of protein expression, isopropyl β-D-1-thiogalactopyranoside (IPTG) was added to a final concentration of 0.5 mM, and the culture was incubated for 3 h at 37°C with shaking at 170 rpm. The collected cell pellet was resuspended in sonication buffer (50 mM Tris-HCl, pH 8.0, 150 mM NaCl, protease inhibitor (167-26101, Wako)) and disrupted by sonication. The supernatant after centrifugation was filtered through a 0.45-μm PVDF membrane and purified. For ΔRING Praja1, after sonication, the lysate was boiled for 15 min, centrifuged, and the supernatant was filtered and purified. Purification of this sample was performed using a HisTrap HP column (5 mL; #17524802, Cytiva) on an ÄKTA avant 25 system (Cytiva). The column was equilibrated with 50 mM Tris-HCl (pH 8.0), 150 mM NaCl, and 50 mM imidazole, and the sample was injected and washed. 50 mM Tris-HCl (pH 8.0), 150 mM NaCl, and 500 mM imidazole were used for elution of His-Praja1.

### Gel shift assay

A gel shift assay was performed to analyze the interaction between DNA and Praja1 in vitro. Unless otherwise noted, pcDNA3.1 empty vector was used as DNA and purified His-tagged Praja1 was used as Praja1. DNA at a final concentration of 5 nM and Praja1 were added to the reaction buffer (100 mM HEPES, pH 7.0, 50 mM NaCl) and incubated at 25°C for 30 min. Genomic DNA was purified from *E. coli* using the Wizard® Genomic DNA Purification Kit and used at a final concentration of 50 μg/μL. As negative controls, His-tag peptide (RMP-0004, CosmoBio) or bovine serum albumin was used. In some experiments, the Praja1-DNA complex was dissociated by adding 1% sodium dodecyl sulfate and 1 mM EDTA followed by heating at 85°C for 5 min. After addition of 10× loading buffer, electrophoresis was performed in cold TAE buffer at 100 V for 15–20 min at 4°C. After electrophoresis, the gel was stained with 0.5 μg/mL ethidium bromide for 20 min and detected using a ChemiDoc Touch imaging system (Bio-Rad). To quantify the fraction of DNA shifted by Praja1 binding, the band intensity of free DNA in the absence of Praja1 was set to 1, and the fraction bound was calculated by subtracting the free DNA intensity under each condition.

### *E. coli* survival assay

pET-HisTEV-Praja1 was transformed into *E. coli* (Rosetta 2 (DE3)) according to the manufacturer’s instruction. For protein expression, colonies were picked into MagicMedia (K6803, Invitrogen) and cultured in liquid medium at 37°C for 16 h. Subsequently, the cell density was adjusted to OD_600_ = 0.5, and the cells were transferred to a quartz cuvette with a 1-mm path length and irradiated with UV-B (254 nm) using a CL1000 UV crosslinker. 1.5 μL of culture were spotted onto agar plates containing ampicillin. Images were captured after culturing for 36 h.

### Cell culture and DNA damage induction

Human neuroblastoma SH-SY5Y cells were cultured in low glucose (1,000 mg/L) Dulbecco’s modified Eagle’s medium (Wako, 041-29775) supplemented with 10% fetal bovine serum (Corning, 35-079-CV) and 1% penicillin-streptomycin (Wako, 168-23191) at 37°C under 5% CO_2_. To induce DNA damage, cells were treated with etoposide (Tokyo Chemical Industry, E0675) at 5 μM for 9 h, KU-55933 (Cayman Chemical Company) at 5 μM for 24 h, or bleomycin sulfate (Cayman Chemical Company) at 7.5 μg/mL for 24 h.

### Transfection

The plasmids used in this study for overexpression of Flag-tagged Praja1 and for expression of Praja1 lacking the nuclear localization signal (ΔNLS-Praja1) were the same as those used in our previous report [12]. EGFP-fused Praja1 was constructed from pRP[Exp]-EGFP/Neo-CAG>Rluc (VectorBuilder) and the Flag-Praja1 expression plasmid using the primers listed below and HiFi DNA Assembly (New England Biolabs, E2621). For transfection of plasmid DNA into SH-SY5Y cells, PEI Max® Reagent (Polysciences, Inc.) or Amaxa™ Cell Line Nucleofector™ Kit V (LONZA, VCA-1003) was used. For siRNA-mediated gene knockdown in SH-SY5Y cells, Lipofectamine RNAiMAX Transfection Reagent (Thermo Fisher Scientific, 13778030) was used. The siRNA used as a negative control and the siRNA for Praja1 knockdown were the same as those used in our previous report [9]. Unless otherwise noted, the culture medium was replaced 24 h after transfection of DNA or siRNA, and cells were used for subsequent experiments. All transfections were performed according to the manufacturer’s recommended protocols.

### Cell viability / Cytotoxicity assay

Cell viability was evaluated using the Viability/Cytotoxicity Multiplex Assay Kit (Dojindo, 346-09271). To induce DNA damage, cells were either exposed to agents or irradiated with 50 mJ/cm^2^ of UV-B using a CL-1000 Ultraviolet Crosslinker (Krackeler Scientific Inc.). Each experiment and the calculation of cell damage rates and cell viability were performed according to the manufacturer’s recommended protocol.

### Comet assay

To visualize DNA damage in cells, a comet Assay was performed using OxiSelect Comet Assay Kits (Cell Biolabs Inc., STA-350) under neutral conditions electrophoresis. Cells were exposed to DNA damaging agents or irradiated with 50 mJ/cm^2^ of UV-B using a CL-1000 Ultraviolet Crosslinker to induce DNA damage. The experiment was conducted according to the manufacturer’s instruction. Images were captured using an inverted microscope IX71 (Olympus). Tail DNA % was calculated by analyzing the captured images using Image J.

### Proteomics analysis

The cells were treated with RIPA Buffer (Wako, 188-02453) supplemented with cOmplete Protease Inhibitor Cocktail (Roche, 4693116001). The lysate was centrifuged at 15,000 rpm at 4°C for 5 minutes, and the supernatant was used as the sample for LC-MS/MS. LC-MS/MS was outsourced to Proteobiologics Co., Ltd. Excessive protein expression, as represented by skin-derived proteins, was excluded as experimental contamination. The preprocessed protein expression list was used for MA plotting and was used as input to the Reactome database [15] to identify varied biological pathways between conditions.

### Statistical analysis

Unless otherwise noted, experiments were repeated at least three times. Error bars in the figures represent standard deviation. Statistical analyses were performed using GraphPad Prism 8. For comparisons between two independent groups, statistical significance was calculated using Student’s *t*-test. For comparisons among three or more independent groups, statistical significance was calculated using one-way analysis of variance (ANOVA). *: P<0.05, **: P<0.01, ***: P<0.001, ****: P<0.0001 (One-way ANOVA post-hoc Tukey’s multiple comparison).

## Results

### Nuclear-localized Praja1 suppresses DNA damage

Praja1 acquired the NLS during mammalian evolution, enabling nuclear localization, but its role in the nucleus remains unclear [12]. To elucidate the function of Praja1 in the nucleus, we transiently expressed wild-type Praja1 and ΔNLS Praja1 in human neuroblastoma SH-SY5Y cells and performed proteomic analysis. The results showed that cells expressing ΔNLS Praja1 exhibited increased expression levels of marker proteins in the DNA damage response pathway (H2A.X, H4C1, POLH, RNF4, UBA52) compared to cells expressing wild-type Praja1 (Figure 1A). These findings suggested that Praja1 plays a functional role against DNA damage. To test this hypothesis, we performed comet assays under neutral conditions on Praja1-expressing cells using three different DNA-damaging agents with distinct mechanisms of action: etoposide, bleomycin, and KU-55933 (Figure 1B). Etoposide acts as a topoisomerase II inhibitor, bleomycin as a DNA strand-breaking agent, and KU-55933 as an inhibitor of the DNA repair factor ATM. The results showed that cells expressing Praja1 exhibited statistically significantly smaller tail DNA % with all three agents, suggesting suppression of DNA damage (Figure 1C-E). ΔNLS Praja1 showed no suppressive effect, indicating that nuclear localization of Praja1 is necessary for this protective effect.

**Figure 1.**
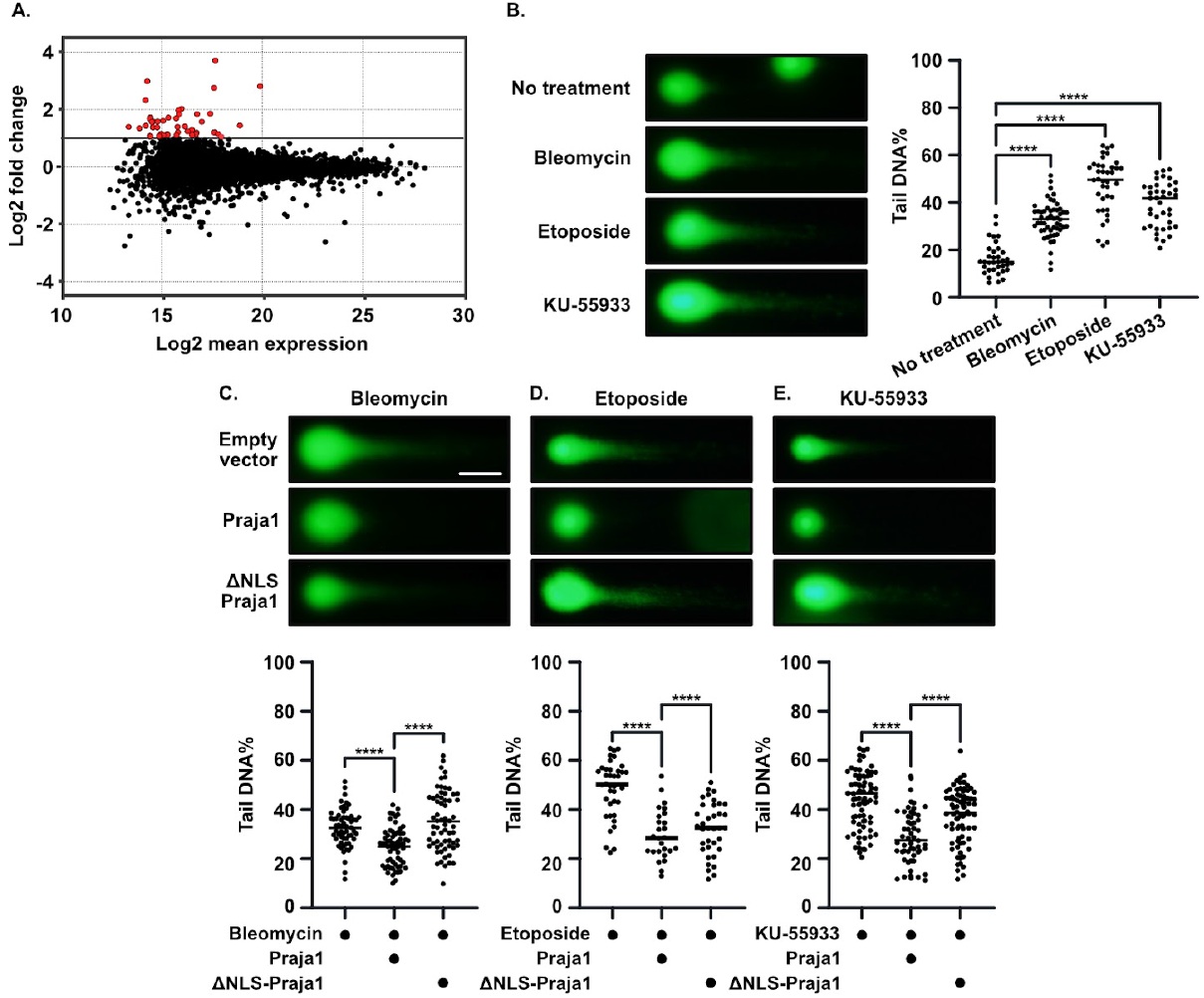
A: Proteomic analysis of Praja1-expressing cells and NLS-deleted PJA1-expressing cells (n=1). Red dots indicate points with Log FC ≥1. B: DNA damage in neutral comet assays upon addition of DNA-damaging agents. SH-SY5Y cells were treated with Bleomycin at 7.5 µg/mL for 24 hours, Etoposide at 5 µM for 9 hours, or KU-55933 at 5 µM for 24 hours. Representative images for each condition are shown. At least 50 comets were quantified by tail DNA % per condition (n=3). ****: P<0.0001 (One-way ANOVA post-hoc Dunnett’s multiple comparison). C–E: Effect of Praja1 on DNA damage in neutral comet assays upon addition of DNA-damaging agents. Representative images for each condition are shown. At least 50 comets were quantified by tail DNA % per condition (n=3). C) Praja1- or ΔNLS Praja1-expressing cells were treated with Bleomycin at 7.5 µg/mL for 24 hours. D) Praja1- or ΔNLS Praja1-expressing cells were treated with Etoposide at 5 µM for 9 hours. E) Praja1- or ΔNLS Praja1-expressing cells were treated with KU-55933 at 5 µM for 24 hours. Scale bar = 20 µm.

### Praja1 knockdown cells exhibit increased sensitivity to DNA damage

We investigated whether endogenous Praja1 can suppress DNA damage-induced cell death. We knocked down Praja1 and evaluated cell viability and cell death following DNA damage induction. Of the three agents mentioned above, KU-55933 was excluded from this evaluation because we could not reproduce the decrease in SH-SY5Y cell viability reported in a previous study [16]. Instead, we added UV-B, which forms cyclobutane pyrimidine dimers in DNA, as an alternative damage source. The results showed that knockdown reduced cell viability and increased cell death in all cases, indicating enhanced sensitivity to DNA damage (Figure 2A-C). These findings suggest that endogenous Praja1 exerts a cytoprotective effect by suppressing DNA damage. Notably, Praja1 knockdown conditions showed a trend of increase in cell viability compared to controls which is consistent with previous studies linking Praja1 to negative regulation of the cell cycle [3]. Rather, the increase in DNA damage-induced cell death despite the cells being in a more proliferative state further supports the protective role of Praja1 in the DNA damage response.

**Figure 2.**
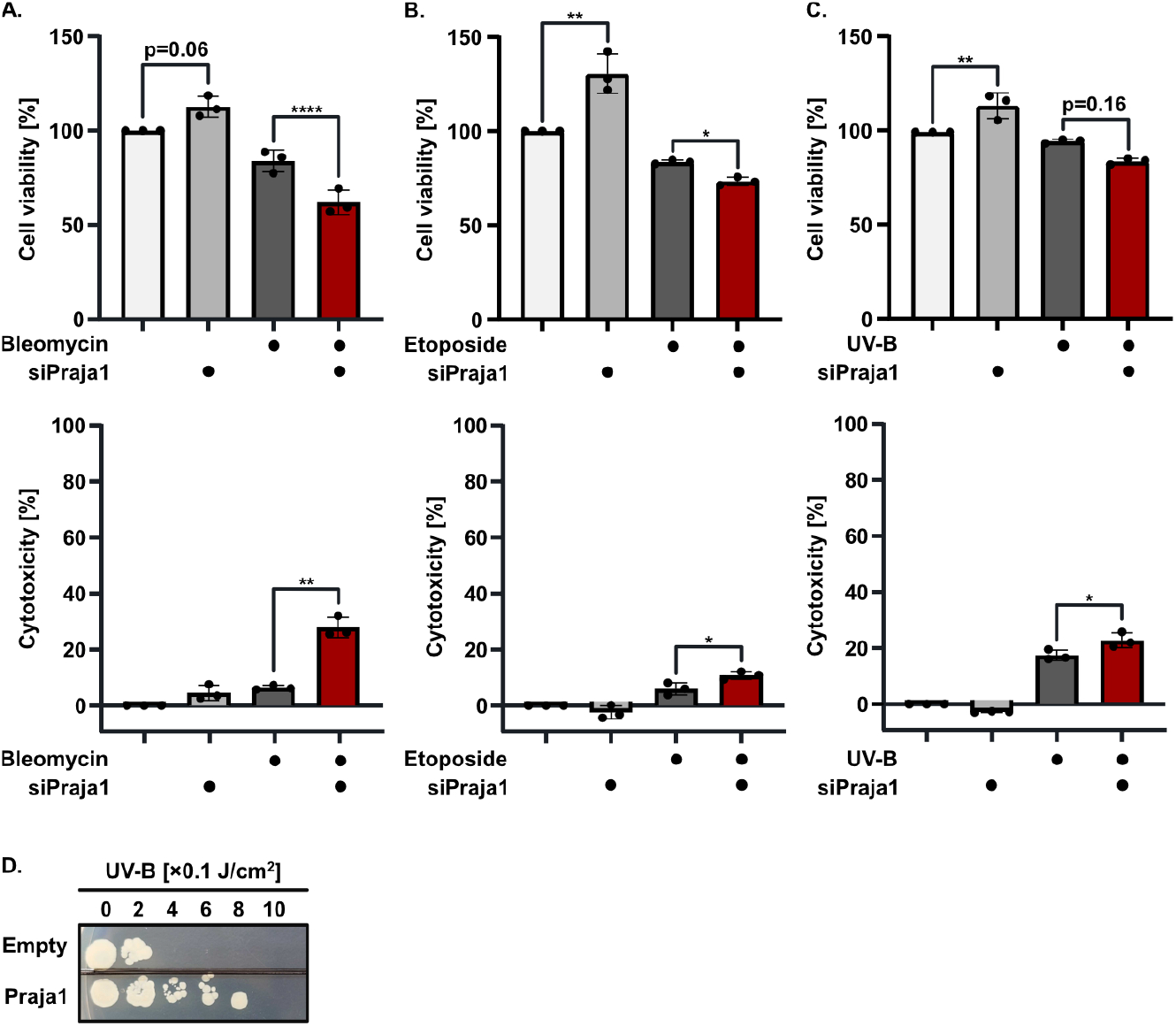
A-C: Effect of Praja1 on cell viability and cytotoxicity upon DNA damaging agent addition. Mean values of data acquired in triplicate were used for each condition. A) Praja1 knockdown cells were treated with Bleomycin at 7.5 µg/mL for 24 hours (n=3). B) Praja1 knockdown cells were treated with Etoposide at 5 µM for 9 hours (n=3). C) Praja1 knockdown cells were irradiated with 50 mJ/cm^2^ UV and subjected to assays 16 hours post-irradiation (n=3). D: Effect on colony formation upon UV irradiation in E. coli. Rosetta2 strain was transformed with an empty vector or Praja1-encoding vector and UV-irradiated during liquid culture (n=3).

Next, we investigated whether the DNA-protective effect of Praja1 requires cooperative molecules. Because Praja1 arose through gene duplication associated with placental mammal divergence [12], *E. coli* lacks endogenous Praja1-interacting molecules. Taking advantage of this, we expressed His-Praja1 in *E. coli* and examined whether DNA damage-induced cell death could be suppressed using a colony survival assay. *E. coli* expressing Praja1 exhibited higher UV irradiation resistance compared to *E. coli* transformed with empty vector (Figure 2D). These results using *E. coli* suggest that Praja1 can exert a DNA-protective effect solely.

### Purified Praja1 directly interacts with DNA

Since Praja1 showed a DNA-protective effect without cooperative molecules, we hypothesized that the molecular basis involves Praja1-DNA interaction. We therefore examined whether purified His-Praja1 binds to the pcDNA 3.1 plasmid vector using a gel shift assay. While no DNA shift was observed in lanes with BSA or His-tag peptide used as negative controls, band shifts were observed in lanes with added Praja1 (Figure 3A), suggesting direct interaction between Praja1 and DNA. We investigated the binding strength and binding mode of Praja1 to DNA by utilizing the principle that binding affinity can be estimated by quantifying gel shift assay results [17]. Since DNA gradually shifted toward higher molecular weight species in a Praja1 concentration-dependent manner, we hypothesized that multiple Praja1 molecules bind to DNA. When the amount of free DNA was fitted to the Hill equation, which assumes interaction of multiple molecules with a single molecule, the IC_50_ was found to be 213 nM (Figure 3B). The Hill coefficient of 2.21 suggests positive cooperativity in this interaction.

**Figure 3.**
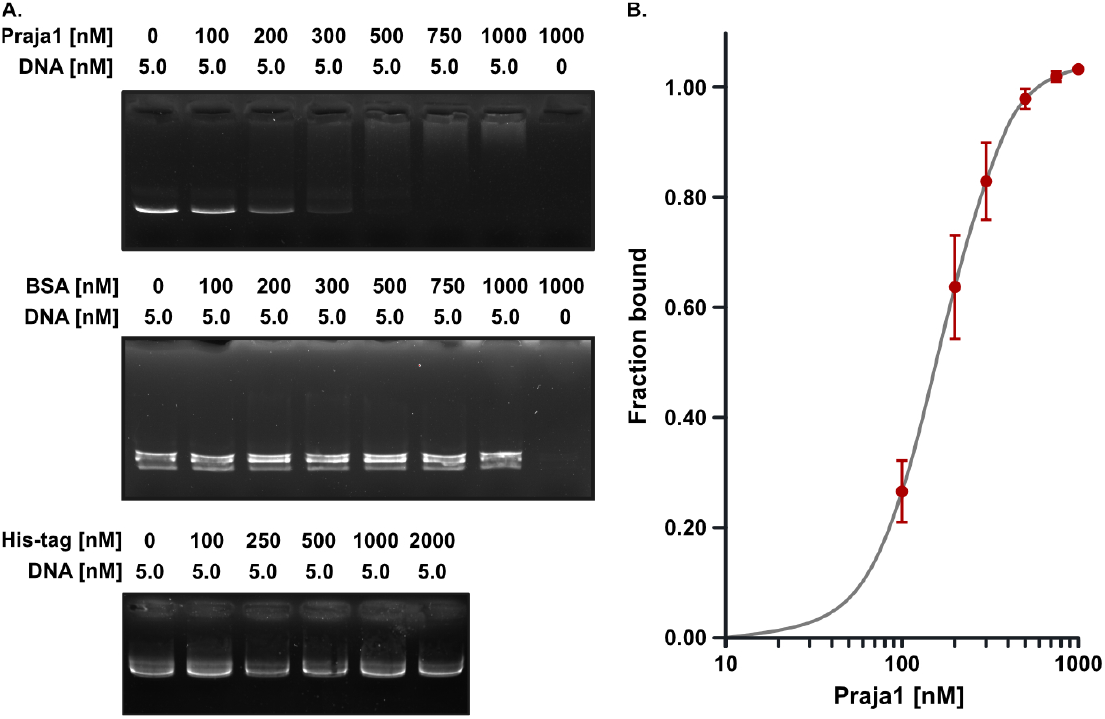
A, B: Gel shift assay of purified Praja1 and pcDNA3.1 plasmid DNA. A) Praja1, bovine serum albumin, or His-tag peptide were mixed with pcDNA3.1 to a total volume of 20 µL, incubated at 25°C for 30 min, and band shifts were observed by agarose gel electrophoresis (n=5). B) Quantification of free pcDNA3.1 was plotted.

### The intrinsically disordered region of Praja1 interacts electrostatically with DNA

Next, we investigated the mechanism driving the Praja1-DNA interaction. Although Praja1 lacks known DNA-binding motifs or structures, it has a high content (28.5%) of amino acids with charged side chains, of which positively charged amino acids that can potentially interact with DNA account for 14.7% of the total (Figure 4A). Furthermore, beside the RING domain, the entire region of Praja1 is intrinsically disordered. Since DNA binding protein tardigrade Dsup has similar properties as Praja1 and interacts electrostatically with DNA [18], we hypothesized that Praja1 also binds to DNA through a similar mechanism. We therefore added salts that shield electrostatic interactions at the time of mixing Praja1 and DNA. The results showed that Praja1-DNA complexes were observed under physiological concentrations of sodium and magnesium found in cells, but complex formation was prevented under excessive salt conditions (Figure 4B). Similarly, no complex was observed when anionic sodium dodecyl sulfate was added. To identify the region responsible for DNA interaction, we focused on the zinc finger, which possesses DNA-binding ability [19,20]. We purified ΔRING Praja1 lacking the RING domain containing the zinc finger and performed the same experiment (Figure 4C). The results showed that ΔRING Praja1 could still interact with DNA, suggesting that the interaction with DNA is driven by the intrinsically disordered region rich in charged amino acids (Figure 4D).

**Figure 4.**
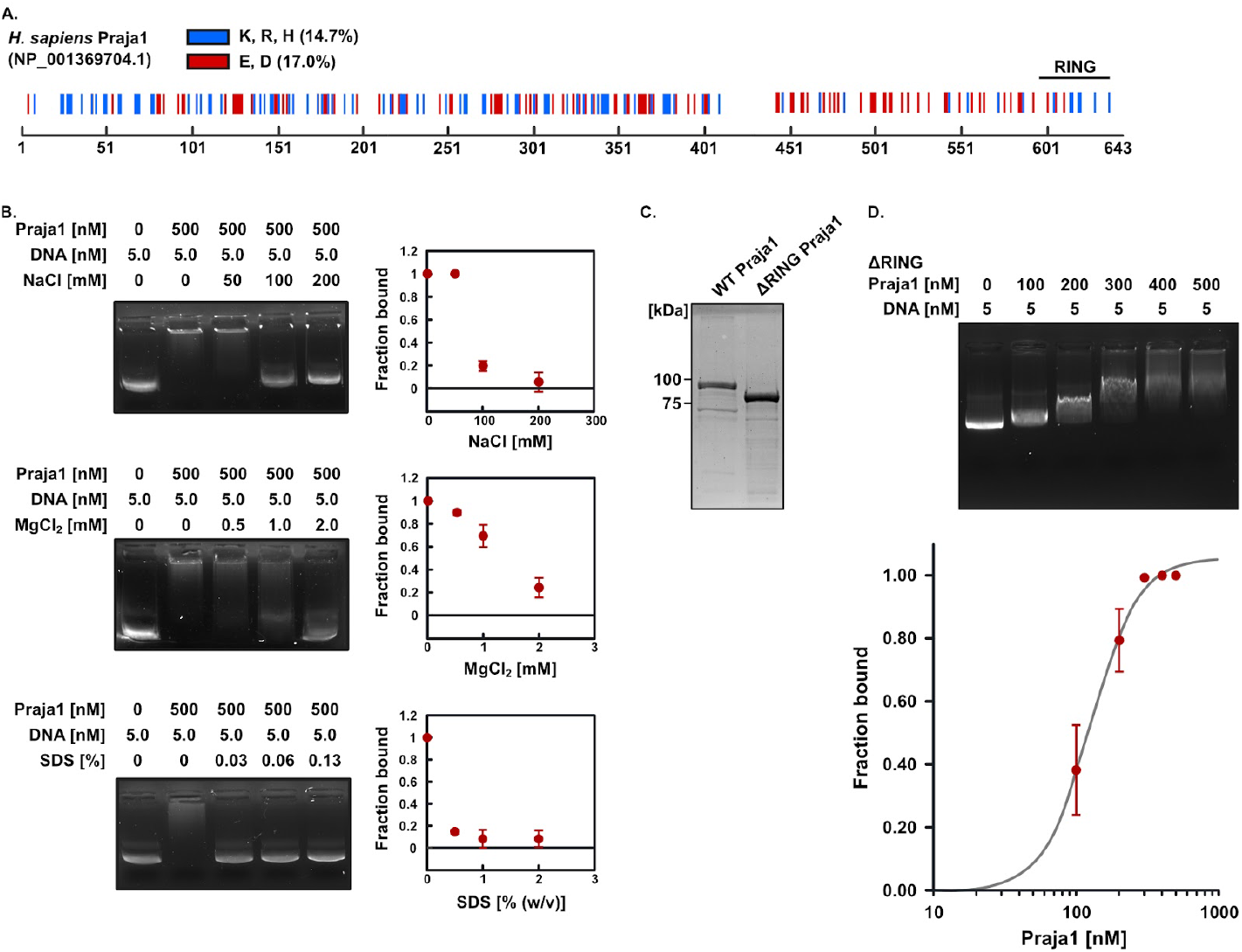
A: Location of charged amino acids and RING domain in H. sapiens Praja1 (GenBank ID: NP_001369704.1). B: Effect of reagents that blocks electrostatic interactions (NaCl, MgCl2, sodium dodecyl sulfate) on Praja1-DNA interaction. Each reagent was mixed simultaneously when combining Praja1 and DNA to a total volume of 20 µL, incubated at 25°C for 30 min, and band shifts were observed by agarose gel electrophoresis (n=3). Mean values from triplicate were plotted. C: RING domain-deleted mutant (ΔRING Praja1) was purified from *E*.*coli*, and molecular weight and purity were confirmed by SDS-PAGE (n=3). D: Gel shift assay of purified ΔRING Praja1 and pcDNA3.1 plasmid DNA. ΔRING Praja1 and pcDNA3.1 were mixed to a total volume of 20 µL, incubated at 25°C for 30 min, and band shifts were observed by agarose gel electrophoresis (n=3). The lower panel shows quantification of free pcDNA3.1.

### Praja1 enhances accessibility of repair machinery to damage sites by altering DNA structure

The molecular basis by which the DNA-protective effect observed in cells is mediated by Praja1 interactions remains unclear. We hypothesized that DNA bound to Praja1 could reduce the effects of damaging agents by Praja1 playing as a protective agent. Gel shift assays were performed by treating Praja1-DNA complexes with DNase I and bleomycin. Contrary to our expectations, DNA degradation was enhanced when DNA is bound to Praja1 (Figure 5A,B). To account for molecular size, we purified genomic DNA from *E. coli* and repeated the experiment, but similarly observed Praja1 binding-dependent DNA degradation (Figure 5C). These may indicate that Praja1 promotes DNA repair efficiency in cells by altering the higher-order structure of DNA.

**Figure 5.**
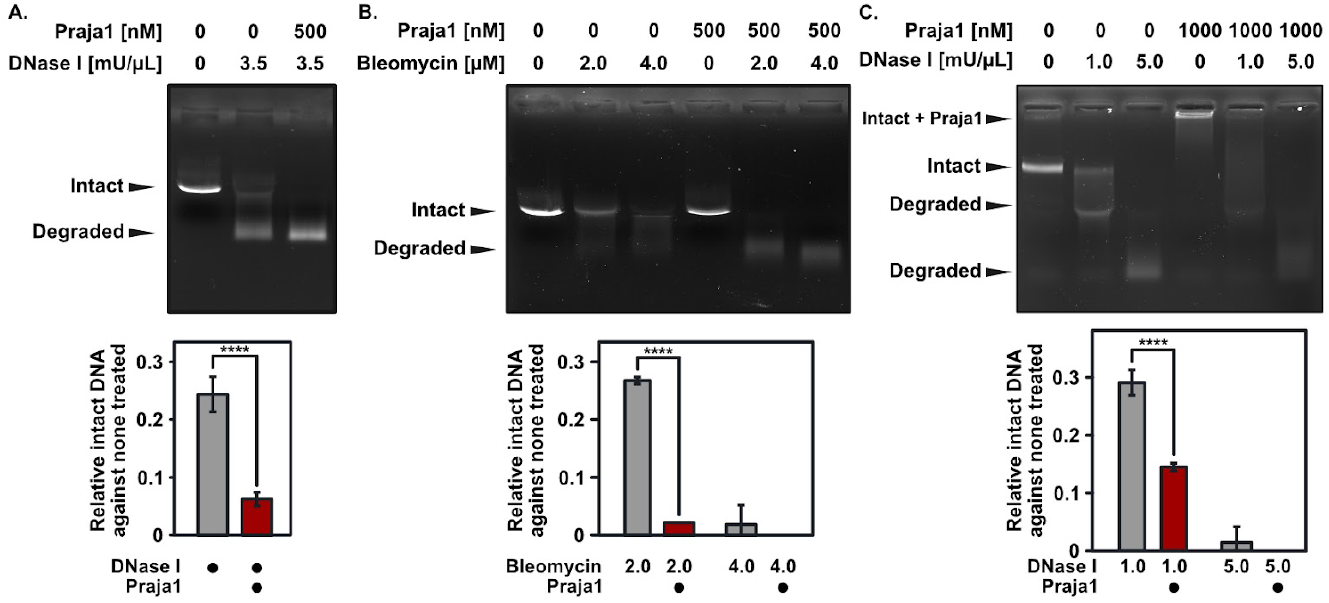
A, B: Gel shift assay was performed on DNA-damaging agents treated Praja1-DNA complexes (n=3). Praja1 and pcDNA3.1 were mixed to a total volume of 20 µL, incubated at 25°C for 30 min, then mixed with A) bleomycin or B) DNase I at 37°C for 15 min. To dissociate DNA from Praja1, 1% SDS and 1mM EDTA were added, followed by heating at 85°C for 5 min. Free pcDNA3.1 not interacting with Praja1 in each lane was quantified (n=3)8. C: Gel shift assay of Praja1 and genomic DNA purified from *E. coli*. Praja1 and genomic DNA were mixed to a total volume of 20 µL, incubated at 25°C for 30 min, then incubated with DNase I at 37°C for 15 min. Band shifts were subsequently observed by electrophoresis (n=3).

## Discussion

In this study, we demonstrated that nuclear Praja1 functions as a direct DNA-binding partner. This property was initially suggested by comparative proteomic analysis between wild-type Praja1 and ΔNLS Praja1, which revealed upregulation of the DNA damage response pathway-related proteins in ΔNLS Praja1-expressing cells. Comet assays demonstrated that Praja1 suppressed the effect of DNA damaging agents, suggesting its DNA-protective effect. Similarly, endogenous knockdown of Praja1 in human neuroblastoma SH-SY5Y cells increased cell death induced by multiple DNA-damaging agents. Furthermore, heterologous expression of Praja1 in *E. coli* enhanced survival under UV irradiation, demonstrating the sufficiency of Praja1 for DNA protection. To understand the molecular mechanism, we first employed gel shift assays to demonstrate electrostatic interactions between the intrinsically disordered region of Praja1 and DNA. Surprisingly, we found Praja1-dependent acceleration of DNA damage in vitro, suggesting structural alterations of DNA, which may, in cells, facilitate efficient access of DNA repair machinery to damaged sites. Collectively, our findings reveal an E3 ubiquitin ligase-independent DNA-protective function of nuclear Praja1 that was acquired during mammalian evolution.

Among the many proteins reported to interact with genomic DNA [21-26], Praja1 is distinguished by its ATP-independent, sequence-nonspecific DNA binding activity as an intrinsically disordered protein. Topoisomerases, one of the most extensively studied classes of DNA-relaxing proteins, resolve DNA supercoiling and entanglement to support fundamental cellular processes including replication, transcription, recombination, and DNA repair [27-29]. These enzymes operate through rigid structural mechanisms, and certain types require ATP [30-32]. In contrast, high mobility group N (HMGN) proteins, like Praja1, interact with genomic DNA primarily through electrostatic interactions [33]. HMGN recognizes nucleosomes and interacts in a largely sequence-nonspecific manner [34-36]. Moreover, HMGN shares with Praja1 the property of being an intrinsically disordered protein [37]. Given that HMGN structurally recognizes nucleosomes and its DNA-interacting motifs have been characterized, future studies should clarify the functional distinctions between HMGN and Praja1[38].

In this study, we demonstrated that Praja1, which possesses numerous charged amino acids, binds DNA through electrostatic interactions, with the IDR serving as the primary DNA-binding region. Similar to Praja1, intrinsically disordered proteins such as HMGN and Dsup interact with DNA electrostatically through multipoint recognition [23,37,39-42]. By analogy, similar approaches may further elucidate the interaction mode of Praja1. Structural biological analyses or approaches such as alanine scanning mutagenesis or random mutagenesis could be possible efficient approaches. A limitation of the current study is that we have not determined whether Praja1 can interact with nucleosomes, DNA with specific structural features or RNA, and if so, whether such interactions are preferred over naked DNA. Additionally, we note that the sufficiency of the molecular basis for the DNA-protective effect has not been fully established. Direct observation of the Praja1-DNA complex will be needed to resolve the contradiction between protective effect in cells and destructive effect in purified systems. While we explained the DNA-protective effect through direct molecular interactions, Praja1 may also exert this effect indirectly through interactions with DDR pathway molecules playing as signalling hubs, which remains to be investigated. Furthermore, we have not addressed what cellular or physiological functions Praja1 exerts through DNA interaction, such as its contributions to neural development and maturation. Resolving these future perspectives will enable discussion of the evolutionary significance of why Praja1 acquired nuclear localization.

## Data Availability

Any additional data will be made available upon request to the corresponding author.

## Author Contributions Statement

Author contributions: Kotaro Kawasaki (Conceptualization [supporting], Formal analysis [equal], Funding acquisition [equal], Investigation [equal], Methodology [equal], Validation [equal], Visualization [equal], Writing – original draft [supporting]), Asahi Toru (Resources [supporting], Software [lead], Supervision [supporting]), Wataru Onodera (Conceptualization [lead], Formal analysis [equal], Funding acquisition [equal], Investigation [equal], Methodology [equal], Project administration [lead], Resources [lead], Supervision [lead], Validation [equal], Visualization [equal], Writing – original draft [lead]).

